# Helical reconstruction in RELION

**DOI:** 10.1101/095034

**Authors:** Shaoda He, Sjors H.W. Scheres

## Abstract

We describe a new implementation for the reconstruction of helical assemblies in the empirical Bayesian framework of RELION. Our approach calculates optimal linear filters for the 3D reconstruction by embedding helical symmetry operators in Fourier-space, and deals with deviations from perfect helical symmetry through Gaussian-shaped priors on the orientations of individual segments. By incorporating our approach into the standard pipeline for single-particle analysis in RELION, our implementation aims to be easily accessible for non-experienced users. Although our implementation does not solve the problem that grossly incorrect structures can be obtained when the wrong helical symmetry is imposed, we show for four different test cases that it is capable of reconstructing structures to near-atomic resolution.

**Abbreviations:**

CARD: Caspase Activation and Recruitment Domain
EM: Electron Microscopy
FSC: Fourier Shell Correlation
IHRSR: Iterative Helical Real-Space Reconstruction
MAVS: Mitochondrial Antiviral Signaling Protein
TMV: Tobacco Mosaic Virus

## Introduction

The first biological structure to be determined by three-dimensional electron microscopy (3D-EM), the extended tail of the T4 bacteriophage, had helical symmetry (De Rosier and Klug, 1968; DeRosier and Moore, 1970). An important advantage for structure determination of helical objects over asymmetrical particles lies in the observation that a single projection image of a helical specimen will typically contain all necessary information to perform a 3D reconstruction. Moreover, for helical reconstructions the number of parameters to be determined is, in principle, strongly reduced compared to singleparticle analysis. That is because in single-particle analysis one needs to determine the relative orientations for every individual particle projection image, while for helical structures many copies of the repeating, asymmetrical unit have fixed relative orientations. Therefore, once the parameters describing helical symmetry and the orientation of the image of an entire filament with respect to that symmetry have been determined, one can reduce the experimental noise efficiently by averaging over a large number of asymmetrical units.

The determination of helical symmetry and the subsequent calculation of a 3D reconstruction were initially performed in Fourier space. The mathematical description of the Fourier transform of an object with helical symmetry was first proposed by Crick, Cochran and Vand (Cochran et al., 1952), and Klug generalised the theory afterwards (Klug et al., 1958). The initial step is to inspect 2D Fourier diffraction patterns of, sometimes averaged, and preferably long and straight, helical filaments. The helical lattice can be considered as a 2D surface lattice that is curled into a cylinder (also see Figure 1). The curling effect causes a convolution of the Fourier transform of the 2D surface lattice with a cylindrical harmonic called the Bessel function. In the 3D Fourier transform of a helical object, discrete horizontal lines, called layer lines, arise from the periodicity along the helical axis. Through the central slice theorem, the radially oscillating ring-like amplitudes on each plane perpendicular to the Z axis in 3D Fourier space give rise to symmetrical maxima across the meridian on each layer line in the Fourier transforms of 2D projections. The position of the maxima along these lines, combined with the realspace width of the helical object, can be used to infer the 2D lattice parameters in a process called Fourier-Bessel indexing. After the Fourier transform of a helix has been indexed, a 3D reconstruction can be obtained through Fourier inversion (DeRosier and Moore, 1970). For more detailed reviews on Fourier-Bessel analysis, the reader is referred to (Diaz et al., 2010; Stewart, 1988).

**Figure 1.**
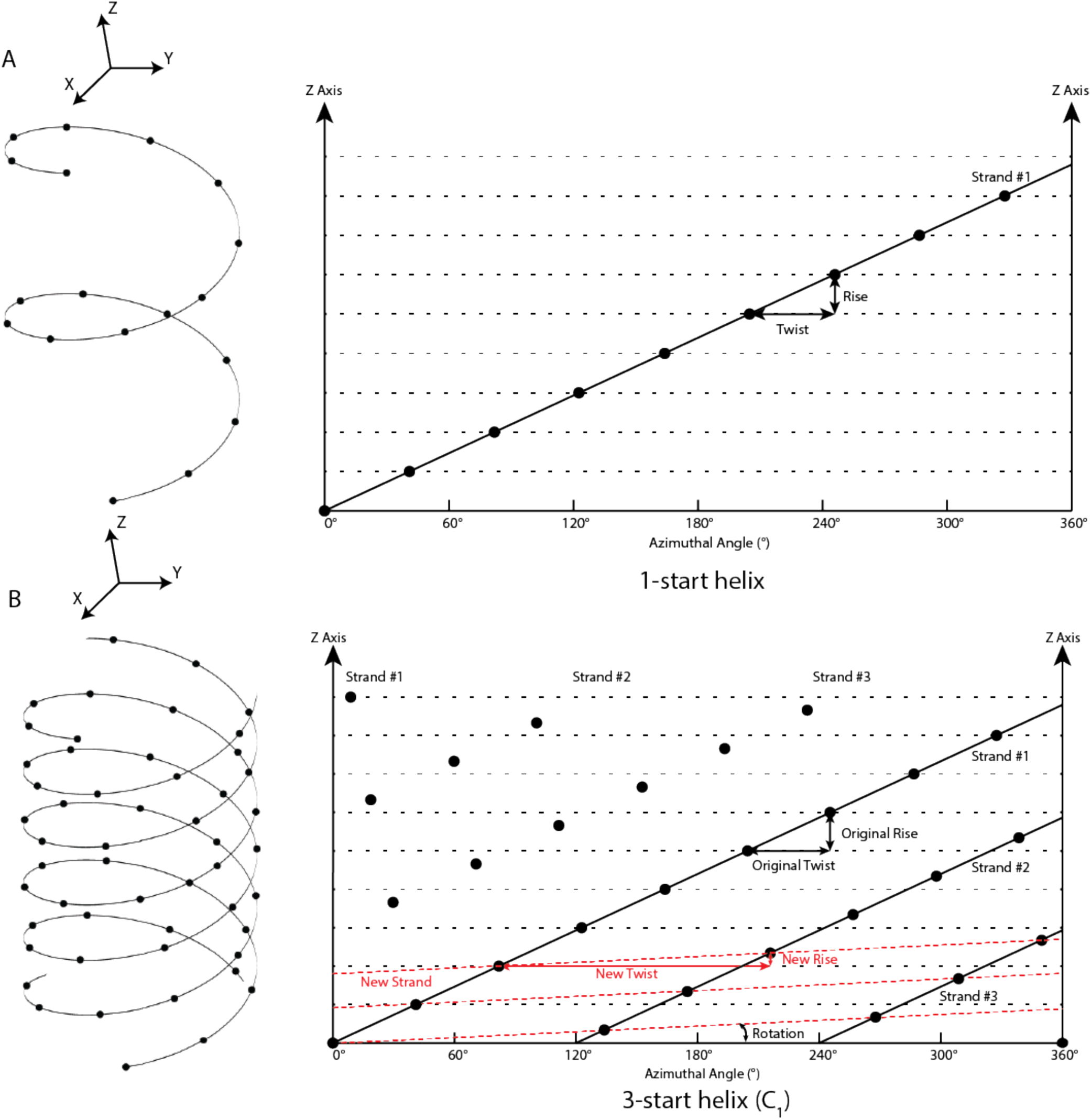
Definition of helical symmetry for *n*-start helices. **A.** Example of a 1-start helix with no point group symmetry (C1). The 3D helical structure on the left can be ‘unfolded’ into a 2D plot of the *z*-height against the azimuthal angle φ on the right. The dots represent the centres of the subunits in the 2D surface lattice. **B)** Example of a 3-start helix (with C1 point group symmetry), where instead of using the original twist and rise of the individual three strands (in black), RELION expresses the helical symmetry using a third of the original rise (in red).

Computer programs for Fourier-Bessel analysis were first developed in the MRC image processing package (Crowther et al., 1996), and have since been adapted in many other packages, such as the Brandeis Helical Package (Owen et al., 1996), Phoelix (Whittaker et al., 1995) and for objects with a break in helical symmetry known as a seam in Ruby-helix (Metlagel et al., 2007). Fourier-Bessel analysis of helices is often considered laborious and difficult, as small inaccuracies in the indexing may lead to incorrect reconstructions. Indexing is particularly prone to errors when the filaments are not lying flat, the lattice is somehow distorted by bending or flattening of the filaments, or by short-range disorder in the helical symmetry. Therefore, the Fourier-Bessel approach only performs well when the specimen adopts a close to perfect helical structure. In practice, many helical assemblies are far from perfect due to molecular flexibility and distortions induced by the sample preparation process. To reduce the effects of long-range distortions on the Fourier-Bessel reconstruction of AChR filaments, Beroukhim and Unwin introduced the idea of dividing helical filaments into short segments, which are independently aligned against a reference structure (Beroukhim and Unwin, 1997).

The introduction of an iterative helical real-space reconstruction (IHRSR) algorithm by Egelman opened up new horizons (Egelman, 2000). Analogous to standard single-particle analysis, the IHRSR algorithm is based on iterative projection matching, where computer-generated projections of a 3D reference in many different orientations are compared with small segments that are cut out from the micrographs in overlapping square boxes along the helical filaments. After the optimal orientations of all segments have been determined in this way, the aligned segments are back-projected into an (asymmetric) 3D reconstruction. The helical twist and rise are then determined by least-square fit within a user-defined search range, and the best symmetry is imposed to generate a symmetric 3D reference for the next iteration. For favourable cases, this method was shown to converge to the correct solution when starting from a solid cylinder (Egelman, 2007), thereby reducing human intervention and enabling helical reconstruction without expertise in Fourier-Bessel analysis. The original IHRSR approach was implemented in the SPIDER package (Frank et al., 1996). More recent implementations were made in FREALIX (Rohou and Grigorieff, 2014) and in SPARX (Behrmann et al., 2012). The SPRING package provides a combination of Fourier-based symmetry analysis and iterative real-space helical processing (Desfosses et al., 2014).

In this paper, we describe an implementation of single-particle-like analysis for helical assemblies in the RELION program (Scheres, 2012a). RELION is based on an empirical Bayesian approach to single-particle analysis in Fourier-space, where a marginalised likelihood function is augmented with a Gaussian prior on the reconstruction (Scheres, 2012b). Optimisation of the corresponding regularised likelihood target is performed using the expectation maximisation algorithm (Dempster et al., 1977). Because parameters of a statistical model for both the signal and the noise are estimated from the data themselves, optimal filters for both the reconstruction and the alignment are determined automatically. Thereby the need for user expertise in devising *ad hoc* filters is strongly reduced. Although helical reconstructions using RELION have been described previously (Clemens et al., 2015; Short et al., 2016; Xu et al., 2015), these approaches relied on external software to impose helical symmetry, so that RELION could not incorporate the additional averaging into its calculation of optimal filters. Moreover, previous versions of RELION were unaware of the signal extending to the edges of the boxed images, and did not exploit statistical priors on the orientations of individual segments to describe deviations from perfect helical symmetry.

Below, in the Methods section, we first describe the modifications to RELION that were made to implement helical reconstruction. Then, in the Results section, we describe how this implementation behaves for four different helical test specimens. We start the Results section with a description of the structure determination procedures for these four test cases, and we describe more detailed analyses of various features of the RELION implementation in the second half of the Results section. Finally, in the Discussion section we analyse the benefits and the remaining pitfalls of our implementation. The approach described in this paper has been implemented in RELION version 2.0, and was accelerated for execution on graphics cards along with the code for standard single-particle analysis of asymmetric particles (Kimanius et al., 2016). RELION is distributed as open-source and can be downloaded for free by both academic and non-academic users from http://www2.mrc-lmb.cam.ac.uk/relion.

## Methods

### Definition of helical symmetry

Helical symmetry in RELION is described by two parameters (Figure 1A): helical *twist* (between −180 and 180 degrees, with positive values corresponding to right-handed helices along the Z-axis) and helical *rise* (> 0 Ångstroms, along the z-axis). Therefore, for any point in real space, the voxel values of a perfectly helical object have the following relationship under cylindrical coordinates with radial distance *r*, angular coordinate *φ*, and height *z* (also see Figure 1A):

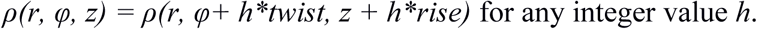

Apart from helical symmetry, additional point group symmetry may also be present. That is, the sample may have an additional *n*-fold rotational symmetry axis along the helical axis (C*n*-symmetry with *n* ≥ 2), and/or it may have a 2-fold rotational axis perpendicular to the symmetry axis (D*n*-symmetry with *n* ≥ 1, where D1 stands for a single 2-fold axis perpendicular to the helical axis but no rotational symmetry along the helical axis). Helices with D*n* symmetry lack polarity, whereas helices with C*n* symmetry typically consist of *n* separate strands of molecules that are packed against each other and that all start at the same level along the z-axis.

In some helical structures the subunit shape and packing may make a particular *n*-start helical family especially prominent, even though the structure does not have C*n* point group symmetry. In such a case the *n* separate helical strands do not all start at the same level along the z-axis. To maintain an intact helical lattice, the difference in translation between the *n* strands have to be multiples of the *rise/n.* Thereby, it becomes possible to describe these so-called *n*-start helical families by a 1-start helix with the following twist and rise (Figure 1B):

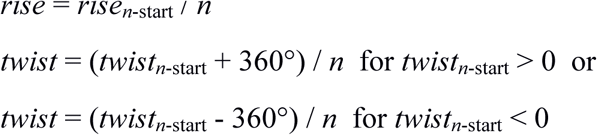

RELION uses the 1-start description of such structures. Although mathematically correct, the 1-start helical lattice may not have an intuitive biological interpretation. The handedness of the 1-start lattice is not necessarily the same as for the *n*-start lattice, and neighbouring molecules do not necessarily contact each other on the 1-start helical lattice.

### Fourier-space symmetrisation and the inter-box distance

In order to estimate the signal-to-noise ratio in the experimental images, which is an intricate part of the empirical Bayesian approach, RELION needs to know over how many asymmetric units a reconstruction is calculated. For asymmetrical particles, each experimental projection contributes a single asymmetric unit to the reconstruction. For helical segments, the amount of overlap between adjacent boxes defines how many new asymmetric units are contained in each new box. Therefore, upon the extraction of helical segments from the micrographs, RELION will ask the user for an interbox distance that is an integer *H* times the helical rise (also see below), so that each new box will contain *H* new asymmetric units. At early stages of processing, when an exact rise may not yet be known, a best guess may be used instead.

For values of *H* larger than one, during the alignment of each segment in the expectation step one could in principle set the 2D Fourier slice *H* times back into the 3D Fourier transform, each time applying a rotation of *h* * *twist* and phase shifts that would correspond to a translation of *h* * *rise* along Z, with *h* = *0,…, H −1* (Figure 2). RELION implements the computationally cheaper, mathematically equivalent option to apply these rotations and phase shifts to the accumulated sum of all slices in 3D Fourier space at the end of every iteration, immediately prior to performing the reconstruction.

**Figure 2:**
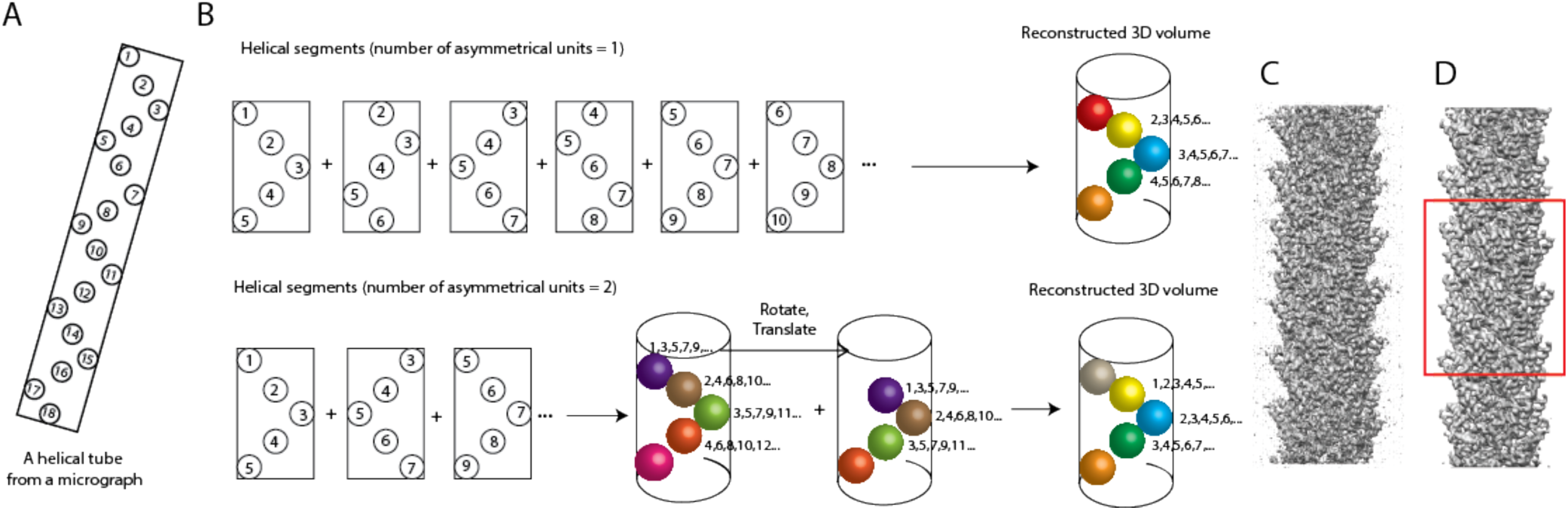
**A.** A schematic representation of a helical filament with 18 asymmetrical units. **B.** Extracted segments with 1 (top) or 2 (bottom) new asymmetrical units per box lead to similar reconstructed volumes after symmetrisation in Fourier-space. Fourier-space symmetrisation is not necessary if the inter-box distance is one helical rise. Otherwise, the reconstructed volume from the helical segments is rotated and translated according to the helical symmetry, and summed up in Fourier space. **C.** The reconstruction of MAVS from manually picked filaments and an inter-box distance corresponding to 6 new asymmetrical units per segment. Without Fourier-space symmetrisation, one does not average over all asymmetrical units. **D.** The MAVS reconstruction after Fourier-space symmetrisation. The central section along the helical axis (within the red box) is averaged over all available asymmetrical units. At the top and the bottom of the helix, blurring is caused by Fourier artefacts (also see main text).

Since the information content of a given set of micrographs is invariant of the inter-box distance, different values of the latter should in principle lead to identical signal-to-noise estimates and 3D reconstructions. Larger inter-box distances, or larger values of *H*, will result in fewer segments and are thus computationally cheaper. However, in practice the useful inter-box distance is limited by artefacts caused by our symmetrisation operations in Fourier-space. For values of *H* larger than 1, translations through phase shifts in the 3D Fourier transform will cause densities at the top of the helix in real-space to "wrap around" to the bottom and *vice versa*. Because the densities at the top and the bottom will typically not coincide, this will lead to additional blurring at the helix ends. Consequently, only the central z-slices in the real-space reconstruction will be intact after the symmetrisation in Fourier space, which places a strict maximum on the inter-box distance of 50% of the box size.

### Real-space symmetrisation and local optimisation of twist and rise

Apart from the artefacts caused by our symmetrisation in Fourier space, other factors will further blur the reconstructed density: helical segments that are not perfectly straight, have varying twist, rise, shrinkage or expansion along the helical axis and inaccuracies in the alignment. Bent helices can be described better by using smaller inter-box distances, which will lead to more segments, and thus more orientational parameters to describe the deviations from a straight helix. But because bent helices can only be approximated with a straight helix in a sufficiently small central section, the top and bottom parts of the reconstructed helix will be more blurred than the central part. The same happens with alignment errors, where the density that is further away from the centre of rotation will be more blurred. As a result, the central part of the reconstructed helix will be better defined than the top and bottom ends. Therefore, it is beneficial to calculate a symmetric structure using only the central z-sections in real-space, and to extend the symmetric structure over the length of the entire box.

As was originally introduced for the IHRSR approach (Egelman, 2000), this real-space symmetrisation can also be accompanied with a local optimisation of the helical twist and rise. This may be necessary, as estimates from traditional Fourier-Bessel indexing or other methods are often not accurate enough for high-resolution reconstruction. Therefore, RELION optionally performs a two-dimensional grid search of the helical twist and rise to find the smallest variance between the averaged sections of the map. Initial search range and step size parameters are provided by the user, and the 2D grid search is iterated with increasingly fine steps around the best twist and rise from the previous iteration. Because the refinement of twist and rise is a local optimisation process, and the energy landscape may contain many local minima, the optimisation of helical parameters should not be considered as a substitute for Fourier-Bessel indexing, asymmetric refinements from featureless cylinders, or experimental approaches like tomography.

### 2D and 3D masks

Masks are used during refinement on both the 2D experimental images and on the 3D reconstruction. Masking the 2D experimental images helps to reduce the influence of noise or the neighbouring particles on the alignment in the expectation step, and the same mask is also used to determine the mean and standard deviation in the background area for the normalisation of boxed segments. Masking the 3D reconstruction reduces the noise in the solvent region, and thus leads to a better reference for the next iteration. For the analysis of helical structures, a circular 2D mask from standard single-particle analysis is multiplied with an additional rectangular mask across the entire particle image with a width according to the *outer_tube_diameter* parameter (Figure 3A). This mask is no longer rotationally invariant, but is rotated according to the most likely in-plane rotation angle for each experimental image. Prior to any refinement, this in-plane rotation is estimated from the manual or template-based particle picking procedures (also see below).

**Figure 3:**
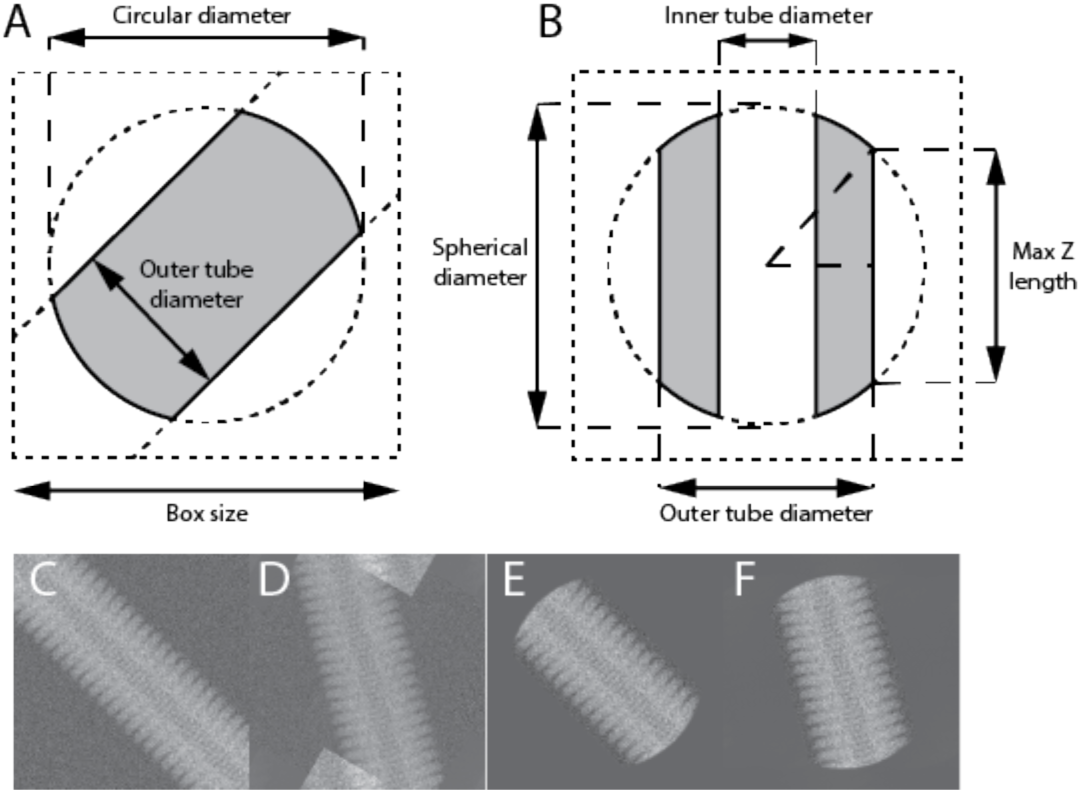
Masks for 2D images (**A**) and 3D maps (**B**). Areas of a filament inside the mask are shown in grey, areas outside the mask in white. In panel A, the helical axis runs at a 45° angle with the horizontal direction, which represents an arbitrary in-plane rotation angle. The helical axis (i.e. the Z-axis) runs parallel to the vertical direction in B. Panels C-F illustrate the need for the circular (or spherical) component of these masks. Rotation of a unmasked segment (**C**) leads to artefacts (top and bottom in **D**), due to wrapping effects caused by rotations in the Fourier domain. The application of a circular mask to the original image (**E**) allows rotation without artefacts (**F**).

Likewise, a spherical solvent mask on the 3D reconstruction from standard single-particle analysis is multiplied with a hollow cylinder mask with the same *outer_tube_diameter* parameter describing the width of the 2D mask. An optional *inner_tube_diameter* parameter also allows to mask away densities inside hollow helices (Figure 3B).

One could argue that because helical segments extend all the way across the image, one would not need the circular or spherical components of these masks. However, as all reference projections are taken as 2D slices from the 3D Fourier transform of the reference map, real-space artefacts may again arise from non-empty corners of the reference upon rotation. To make comparisons between the reference projections and the experimental images consistent, the corners are removed from both the 3D map and the 2D images (Figure 3C-F).

### Incorporating prior information about the orientations

The statistical framework provides an elegant way to express prior information about the relative rotations and translations of each experimental image. For example, in both single-particle analysis and the processing of helical filaments, (manual or automated) particle picking will yield residual translational offsets around the approximate centre of the images. RELION expresses this information as a Gaussian prior probability function, and estimates its standard deviation from the data during the iterative refinement. For helical processing, this distribution is changed into an anisotropic function (Figure 4A). Along the helical axis, the prior probability is a tophat distribution with non-zero values from −*rise*/2 to +*rise*/2 around the centre of the image. This distribution prevents segments from translating into the next asymmetric unit, which would lead to averaging multiple times over the same signal in overlapping segments. In the orthogonal direction, a Gaussian distribution is used, for which the standard deviation is re-estimated at every iteration of the expectation maximisation algorithm in the same way as in the single-particle approach (Scheres, 2012a).

**Figure 4:**
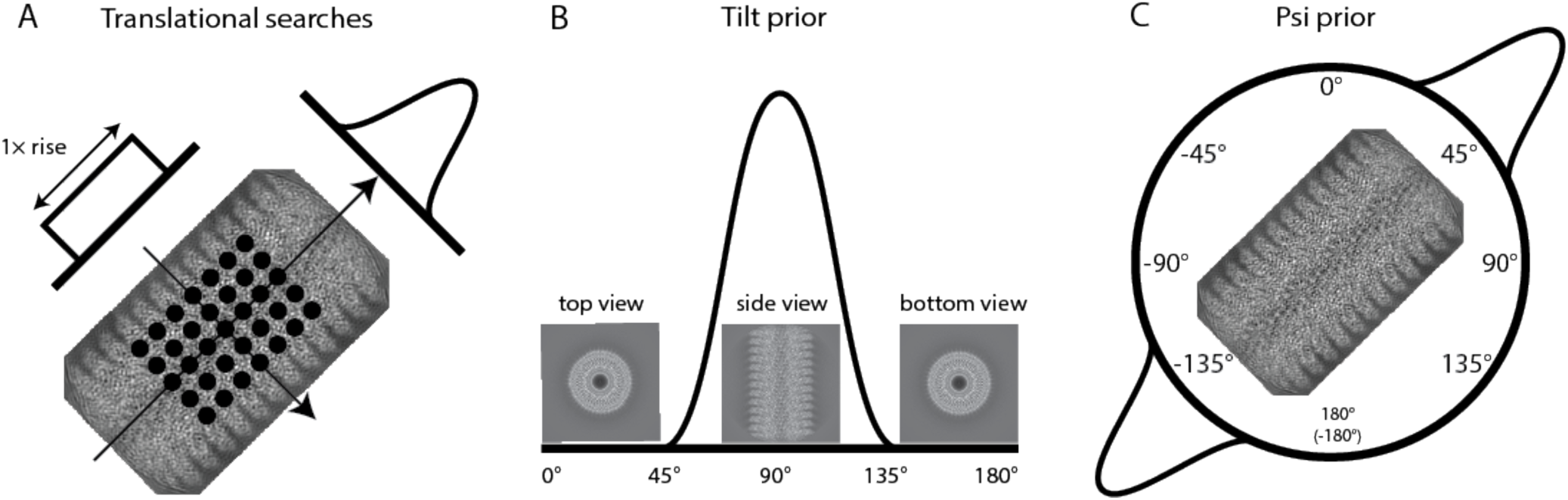
Priors on the orientational parameters. **A.** A top-hat prior on the in-plane translations along the helical axis, and a Gaussian prior on the in-plane translations perpendicular to the helical axis. **B.** A mono-modal Gaussian prior on the tilt angle, which describes the out-of-plane rocking of filaments in the ice layer. **C.** A bi-modal Gaussian prior on the psi angle, which describes the in-plane rotation of the filament.

Unlike in single-particle analysis, there is also ample prior knowledge about the relative rotations of helical segments. Firstly, because the long filaments tend to lie approximately horizontal inside the ice layer, we use a Gaussian distribution that is centred at 90° on the tilt angle in RELION (Figure 4B). Distributions with a wider standard deviation can be used to account for increasing degrees of rocking, but at present this parameter is fixed at a value provided by the user. Alternatively, the user can choose to allow the centre of the tilt prior for each particle to move to the most likely angle from the previous iteration. In principle, if the helical twist and rise are known, there are also strong relationships for neighbouring segments between the rotations along the helical axis, i.e. the ‘rot’ angle in RELION. However, as the rot angle is strongly coupled with the translation along the helical axis, expressing this information into a prior is more complicated, and this is not done in the current implementation.

For the in-plane rotations, the ‘psi’ angle in RELION, we introduce a bimodal prior for helical segments that expresses the prior knowledge that the particle picking provides about the direction of the helix, but with an ambiguity in its polarity (Figure 4C). This prior consists of two opposing Gaussian distributions that are clipped at 3 standard deviations. The ambiguity in polarity is absent for helices with D*n* point group symmetry. For helices with C*n* symmetry, this polarity is typically resolved during the refinement process. One can also express the prior knowledge that all segments from the same helix should have the same polarity. In our implementation, one starts the refinement with symmetrical priors on the in-plane rotations, where the two opposing Gaussian distributions have equal weights. As the refinement progresses, and particles accumulate a stronger probability for one of the two polarities, the difference in the weights on the two opposing Gaussians becomes correspondingly larger. In 3D auto-refinement, and sometimes also in 2D or 3D classification, initial iterations of exhaustive orientational searches are followed by iterations where only local orientational searches are performed. At this point, the local searches are only performed around the in-plane rotation with the largest weight.

For both the in-plane rotation and the tilt-angle, we also implemented an option to centre the prior on the optimal orientations from the previous iteration for all neighbouring segments within a user-defined distance. A Gaussian function (with a standard deviation of one third of the user-defined distance) is used to assign lower weights to segments that are further away. This local averaging option is implemented in the `relion_refine` program as `--helical_sigma_distance X`, where X is the standard deviation of the Gaussian weight function. Local averaging reduces discontinuities in the orientational assignments along a single filament.

### Particle picking and extraction

Apart from modifications to the refinement program that underlies the 2D/3D classification and 3D auto-refinement options of RELION, we also modified the programs for manual and template-based particle picking and particle extraction. The program for manual picking allows the user to identify the start and end points of helices by subsequent left-mouse clicks in the micrographs. For the template-based picking algorithm, the figure-of-merit calculation, which expresses the fit between the template and any local area in the micrograph, remains unchanged from the single-particle approach (Scheres, 2015). The adaptation of the algorithm lies in the detection of filaments and cross-overs between filaments. The latter requires additional parameters for the helix diameter, the inter-box distance (in multiples of the estimated helical rise), and the maximum curvature of the filaments (which is expressed as the fraction, between 0 and 1, of the curvature of a circle with the particle radius). The algorithm to detect filaments searches for peaks in the figure-of-merit map for the entire micrograph, and considers neighbouring peaks within a distance of half the template box size. Peaks that have too few neighbours, or neighbours that do not satisfy the maximum curvature requirement, are discarded. Then, the algorithm iteratively extends candidates for helical tracks in both directions, starting from the peak with the highest figure-of-merit, until no new segments can be at the ends. Once all helical tracks have been identified, the program calculates output segment coordinates according to the specified inter-box distance. At this stage, regions in the micrographs where two different helical filaments overlap are excluded.

Upon extraction of the segments from the micrographs, apart from the corresponding X-Y coordinates and the figure-of-merit, the particle extraction program also outputs for each segment its in-plane rotation along the helical track, a distinct number for the helix it belongs to, and its position within that helix. The latter information is then used to construct the relevant priors on the orientations of each segment as described above. This process does not require any additional user input when RELION is used for all steps from particle picking to refinement. However, if users have previously picked particles in a program outside the RELION workflow, then the additional metadata should be provided through an imported STAR file in RELION format. In that case, it may be more convenient to re-pick and/or re-extract the segments inside RELION.

## Results

### Test data sets

To test the different aspects of our implementation we used four data sets (Table 1). Three of these data sets are available from the EMPIAR data base (Iudin et al., 2016): EMPIAR-10020 on tobacco mosaic virus (TMV) (Fromm et al., 2015); EMPIAR-10019 of the contractile sheath of the bacterial type-VI secretion system, which is composed of the proteins VipA/VipB (Kudryashev et al., 2015); and EMPIAR-10031 on prion-like aggregates of the mitochondrial antiviral signalling protein that are mediated by the N-terminal caspase activation and recruitment domain (MAVS/CARD) (Xu et al., 2015). The fourth data set was collected as part of an in-house research project on the bacterial actin homolog, MamK (Löwe et al., 2016). These test data sets were collected on different direct-electron detectors, while the filaments have different helical and point group symmetries and vary in their levels of order. TMV has been used as a test specimen for other helical reconstruction programs, e.g. (Desfosses et al., 2014; Rohou and Grigorieff, 2014), and consists of straight and regular helical assemblies that are interspersed with disc-like structures that break the helical symmetry. VipA/VipB filaments are also relatively straight and regular, while MAVS/CARD and MamK filaments are more flexible.

**Table 1:**
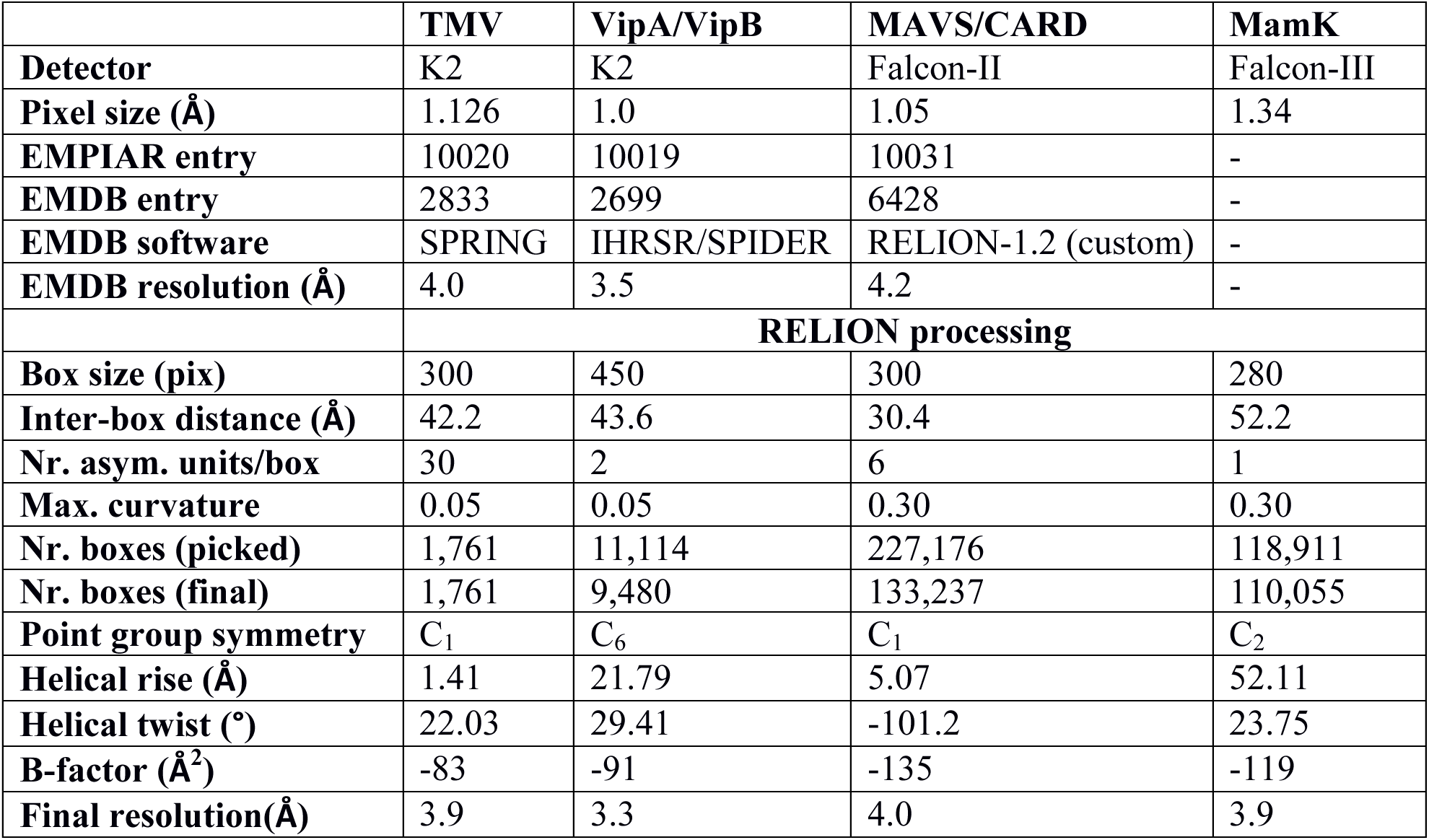
Test data set characteristics

|  | **TMV** | **VipA/VipB** | **MAVS/CARD** | **MamK** |
| --- | --- | --- | --- | --- |
| **Detector** | K2 | K2 | Falcon-II | Falcon-III |
| **Pixel size (Å)** | 1.126 | 1.0 | 1.05 | 1.34 |
| **EMPIAR entry** | 10020 | 10019 | 10031 | - |
| **EMDB entry** | 2833 | 2699 | 6428 | - |
| **EMDB software** | SPRING | IHRSR/SPIDER | RELION-1.2 (custom) | - |
| **EMDB resolution (Å)** | 4.0 | 3.5 | 4.2 | - |
|  | **RELION processing** |  |  |  |
| **Box size (pix)** | 300 | 450 | 300 | 280 |
| **Inter-box distance (Å)** | 42.2 | 43.6 | 30.4 | 52.2 |
| **Nr. asym. units/box** | 30 | 2 | 6 | 1 |
| **Max. curvature** | 0.05 | 0.05 | 0.30 | 0.30 |
| **Nr. boxes (picked)** | 1,761 | 11,114 | 227,176 | 118,911 |
| **Nr. boxes (final)** | 1,761 | 9,480 | 133,237 | 110,055 |
| **Point group symmetry** | C<sub>1</sub> | C<sub>6</sub> | C<sub>1</sub> | C<sub>2</sub> |
| **Helical rise (Å)** | 1.41 | 21.79 | 5.07 | 52.11 |
| **Helical twist (°)** | 22.03 | 29.41 | -101.2 | 23.75 |
| **B-factor (Å<sup>2</sup>)** | -83 | -91 | -135 | -119 |
| **Final resolution(Å)** | 3.9 | 3.3 | 4.0 | 3.9 |

### Particle picking and 2D classification

Figure 5 shows a representative micrograph for each of the four test samples. Because the discs in TMV filaments are hard to distinguish using the template-based picking approach, we performed the picking of TMV filaments manually. The relatively straight VipA/VipB filaments were also picked manually. MAVS/CARD and MamK filaments are increasingly bent. Because the manual picking approach assumes straight filaments, template-based particle picking is more suitable for these cases. The corresponding templates were generated from initial 2D classification runs on segments that were obtained by manually picking of the straighter filaments in a subset of the micrographs.

**Figure 5:**
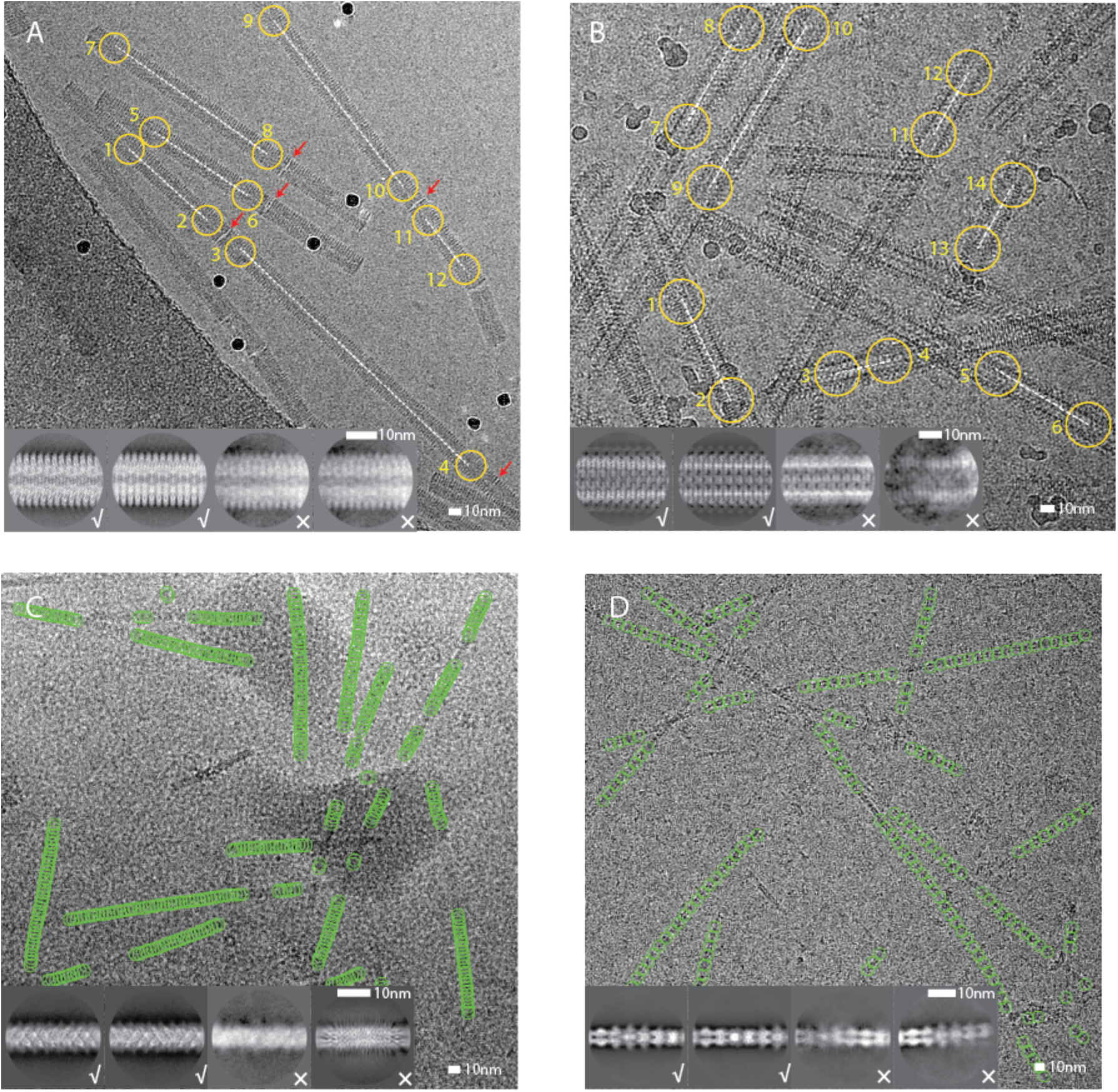
Representative micrographs, particle picking and 2D classification results for the four test cases. **A.** Manually picked coordinates (yellow) for a TMV micrograph. Non-helical discs are indicated with a red arrow. **B.** Manually picked coordinates (yellow) for a VipA/VipB micrograph. **C.** Autopicked coordinates (green) for a MAVS/CARD micrograph. **D.** Auto-picked coordinates (green) for a MamK micrograph. For each test case, four 2D class averages are shown as insets. Good class averages are indicated with a tick, suboptimal ones with a cross.

After manual or template-based particle picking of the entire data sets, all extracted segments were subjected to a single round of reference-free 2D classification, which was used to separate suboptimal segments from segments that contribute to high-resolution class averages (Fig. 5A-D, insets).

For the VipA/VipB and the MAVS/CARD filaments, we also compared how manual and template-based particle picking affected the resolution of the final reconstruction after 3D auto-refinement (Supplementary table 1). For this comparison, we used the manually picked coordinates of MAVS/CARD that were available in the EMPIAR-10031 entry. For both test cases, auto-picking was again performed with 2D class averages from a small subset of manually picked segments. Template-based particle picking performed at least as well as manual particle picking. In particular for the more bent MAVS/CARD filaments, template-based particle picking selected many more particles, which gave rise to a higher resolution map after 3D auto-refinement.

### Initial model generation

Because the expectation maximisation algorithm in RELION is a local optimiser, the initial 3D reference needs to be within the radius of convergence of the optimisation algorithm in order to yield a correct structure. This is also true for alternative IHRSR implementations. In particular, providing incorrect helical twist and rise has been associated with nasty false minima that lead to grossly incorrect structures, even at apparently relatively high resolutions (reviewed by Egelman, 2010, 2014; Sachse et al., 2007). Thereby, initial model generation and the corresponding estimation of helical parameters probably represent the most critical steps in successful helical reconstruction, and the user has multiple options here. Additional experimental data, e.g. from tomograms, sub-tomogram averages, or from homologous structures may be an efficient way to obtain initial 3D models. For example, in the original publication on the MamK structure (Löwe et al., 2016), we placed the atomic coordinates of a crystal structure of the MamK subunit in a helical lattice that was similar to actin, using the `-- pdb_helix` option in the `relion_helix_toolbox` program. Nevertheless, for all four test data sets described here, we chose to calculate initial models *de novo* (Figure 6).

**Figure 6:**
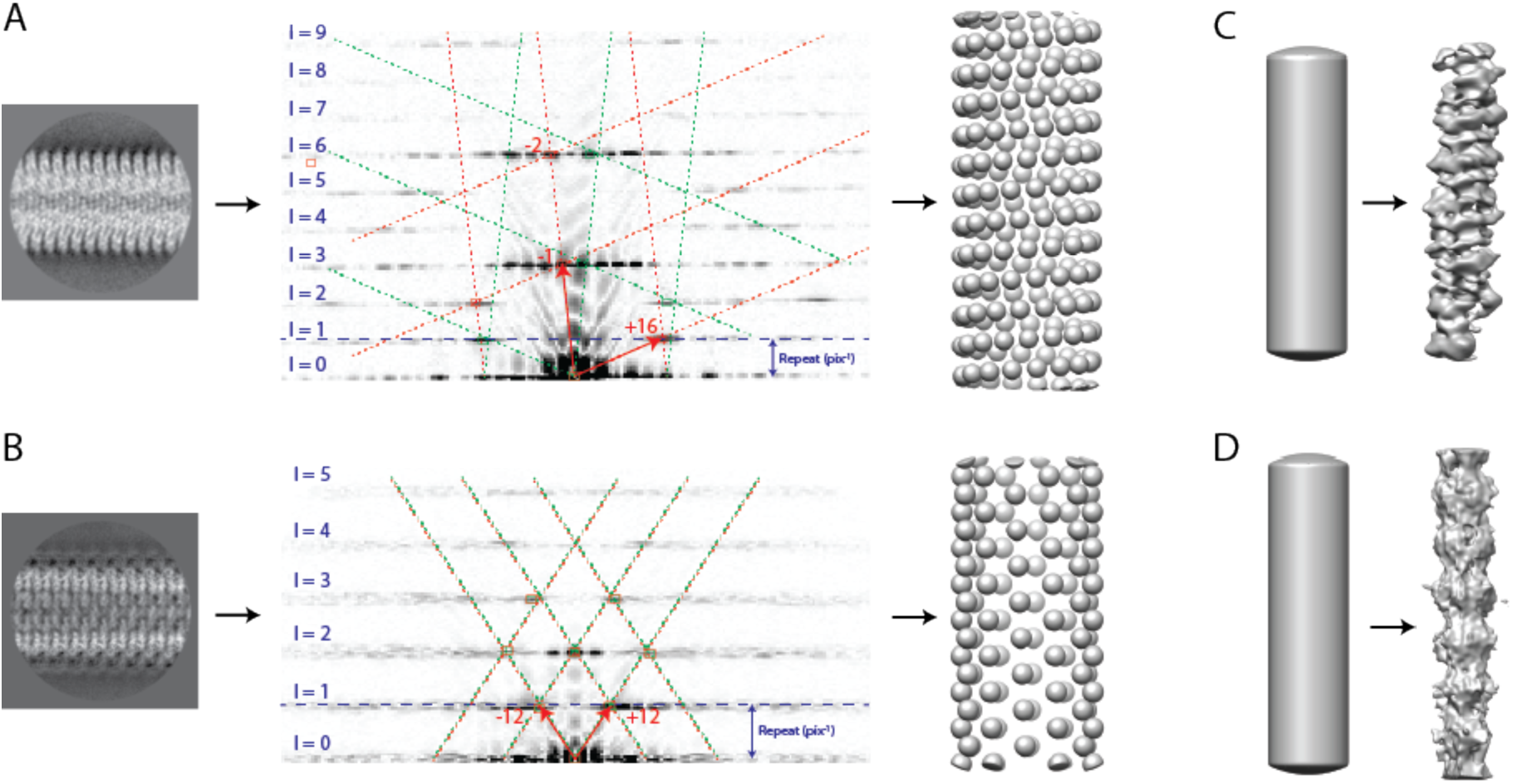
Initial model generation for TMV (**A**), VipA/VipB (**B**), MAVS/CARD (**C**) and MamK (**D**). In A and B, on the left the amplitudes of Fourier-space diffraction pattern of a selected 2D class average image is shown, with the layer lines numbered and the vector pair that constructs the helical lattice indicated with red arrows. The helical lattice and its mirror counterpart are shown with red and green boxes, respectively. The corresponding initial spheres-only models are shown on the right. In C and D, the featureless cylinder (the width of which is estimated from the 2D class averages) that was used as an initial model is shown on the left, and the result from a 3D refinement without imposing any symmetry is shown on the right.

For both TMV and VipA/VipB, we obtained initial estimates for the helical twist and rise from indexing the diffraction pattern of a 2D class average image (Fig 6A,B). In brief, we first calculate the helical repeat distance, which is the distance between two subunits which are (approximately) registered along the z-axis, from the reciprocal distance from the equator to the first layer-line (see blue dashed lines in Fig 6A,B). Then, to determine how many subunits there are in a single turn, we mark the first maxima along each layer-line and find a 2D lattice that links half of these maxima, while the mirrored lattice across the meridian links the other half of the maxima. This mirror symmetry can be thought of as representing two overlapping lattices from the near and far sides of the cylindrical structure (shown with red and green boxes in Fig 6A,B). For the two unit vectors that span this lattice, we then estimate the order of their Bessel functions from their distances to the meridian and the realspace diameter of the helix as observed in the 2D class averages. The Bessel orders and the geometry of the lattice in reciprocal space, are then used to calculate the helical pattern in real-space (Supplementary Figure 1), from which the helical twist and rise can be derived.

For TMV, the unit vectors of the lattice on layer lines 1 and 3, have estimated Bessel orders +16 and −1, respectively. This leads to 16⅓ subunits in a full turn in real-space, and hence an estimated twist of 22.04°. The 16⅓ subunits per turn results in the same structure being in register along the z-axis after 3 turns. Therefore, the estimated repeat distance of 71 Å is divided by 3×16⅓=49 to yield an estimated rise of 1.45 Å. For VipA/VipB, the unit vectors of the lattice both lie on the first lattice line, and have estimated orders of +12. so initial estimates for twist and rise were 30° and 21.4 Å. Initial 3D models were then generated for both test cases by placing spheres in a helical pattern with the estimated twist and rise. The latter functionality has been implemented through the `--simulate_helix` option in the `relion_helix_toolbox` program, where the user also provides the diameter of each sphere (so that neighbouring spheres touch each other to form the helical lattice) and the diameter of the helix (as obtained from 2D class averages). Subsequent refinement of these multi-sphere models, while imposing helical symmetry and performing local optimisations of the helical twist and rise, then led to correctly converged structures at near-atomic resolution for both TMV and VipA/VipB. The final, refined values of twist and rise were 22.03° and 1.410 Å for TMV, and 29.41° and 21.79 Å for VipA/VipB.

For the MAVS/CARD and the MamK helices, we started 3D refinements from a featureless cylinder. The `relion_helix_toolbox` program implements functionality to create such models through the `--cylinder` option, where the user only needs to provide its diameter. For both test cases, refining a cylindrical model in C1, without imposing any helical symmetry (using the `-- ignore_helical_symmetry` option in the `relion_refine` program), yielded a map in which helical symmetry was already apparent. The `--search` option in the `relion_helix_toolbox` program was then used to estimate the helical twist and rise from these asymmetrical reconstructions. This yielded estimates of −101.1° and 5.150 Å for MAVS/CARD and 18.29° and 55.24 Å for MamK. Using a symmetrised version of the asymmetrical map, while again imposing helical symmetry and performing local optimisations of the helical twist and rise, then also led to correct convergence. The final, refined values of twist and rise were −101.2° and 5.071 Å for MAVS/CARD, and 23.75° and 52.11 Å for MamK.

### High-resolution structure determination

At this stage of the structure determination process, the workflow of high-resolution structure determination becomes more similar to the standard approach to single-particle analysis, see (Scheres, 2016) for a recent review. For each of the four test cases, we performed an initial 3D auto-refinement with the selected segments. For the TMV and MAVS/CARD data sets, we then extracted movie-frames for the selected particles and performed per-particle beam-induced motion correction and resolution dependent radiation-damage weighting for each movie frame, which is known as particle polishing in the RELION work flow (Scheres, 2014). Although movie frames were available for our MamK data set, and particle polishing was performed when we initially published this data set (Löwe et al., 2016), we chose not to re-process the movie frames of this large data set for this paper. The VipA/VipB EMPIAR entry does not contain movies, so no movie processing was performed.

After 3D auto-refinement of the polished particles, we attempted further selection of the best segments by 3D classification. In these classifications, we did not allow further adjustments of the orientations. However, for none of the four test cases 3D classification managed to identify a subset that yielded a higher resolution map than we had already calculated from all polished particles. Using RELION’s post-processing job-type, we then applied automated B-factor sharpening (Rosenthal and Henderson, 2003) and corrected the FSC between the two independently refined maps of the 3D auto-refinement for the convolution effects of a solvent mask using phase randomisation (Chen et al., 2013). The final overall resolution estimate was based on the FSC=0.143 threshold (Scheres and Chen, 2012). Variations in local resolution were estimated using phase-randomisation in combination with sliding a soft spherical mask across the entire volume (Fernandez-Leiro and Scheres, 2016a). After postprocessing, we used the `relion_helix_toolbox` program with the `--impose` option to symmetrise the (asymmetric) postprocessed map according to the refined helical twist and rise. Figure 7 shows the final maps, the solvent-corrected FSC curves and the local resolution estimates. Parameters specific to the helical reconstruction, estimated B-factors, overall resolution estimates and other refinement characteristics are summarised in Table 1. 2D classification or 3D auto-refinement for the smaller data sets of TMV and VipA/VipB took a few hours on 64-128 CPU cores. Similar calculations with the larger data sets of MAVS/CARD and MamK took up to an entire day using similar computer resources. A modern GPU (like the Nvidia GTX1080) processes the equivalent of approximately 100 CPU cores in these calculations.

**Figure 7:**
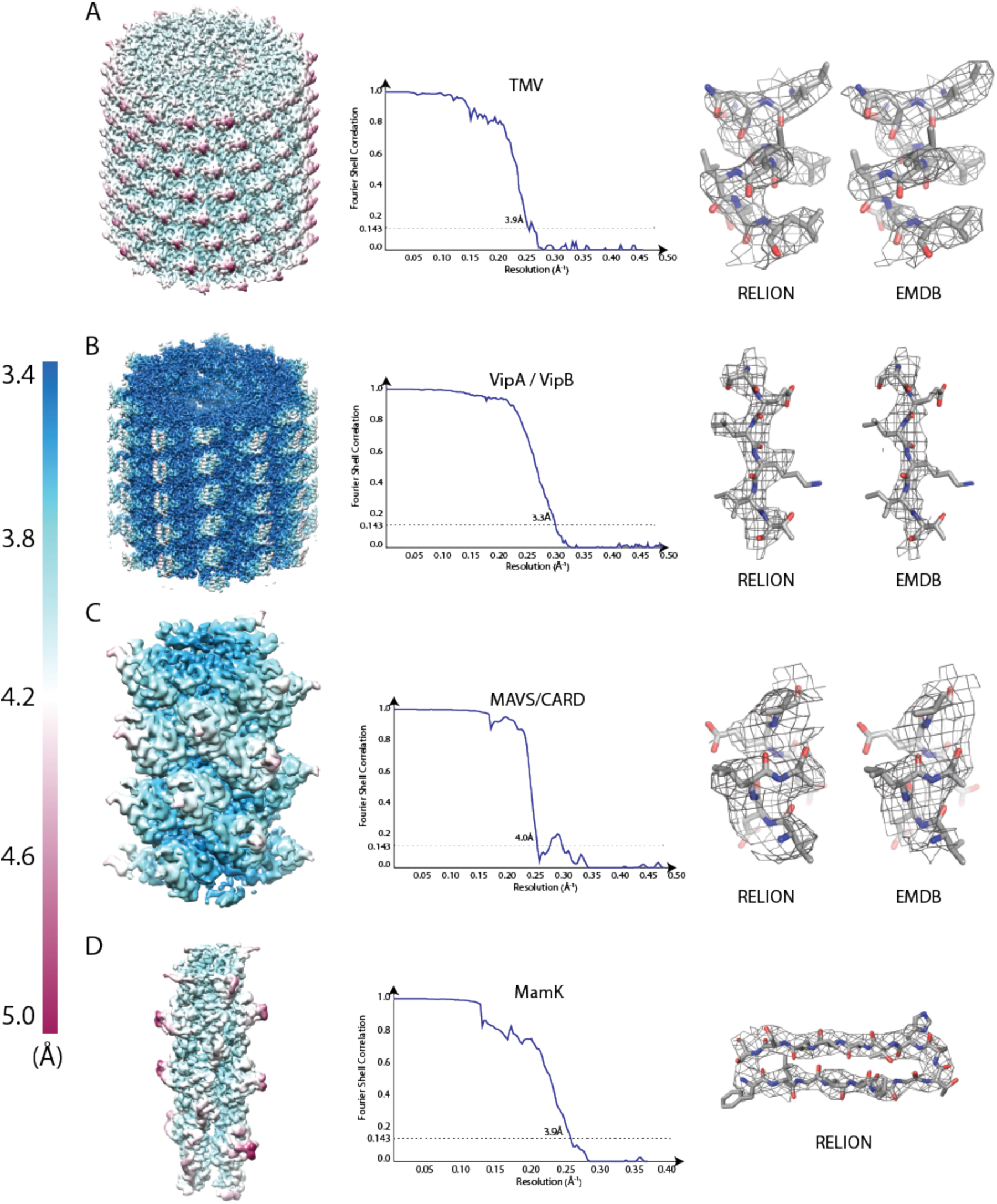
Reconstructions for the four test cases: TMV (**A**), VipA/VipB (**B**), MAVS/CARD (**C**) and MamK (**D**). A view of the reconstructed helix coloured according to local-resolution estimates (from 3.5 Å in blue to 5 Å in maroon) is shown on the left. The FSC curve between the two independently refined half-sets is shown in the middle. A detail of the structure with the corresponding atomic model is shown on the right. For TMV, VipA/VipB and MAVS/CARD, the same detail from the EMDB map is also shown.

### Local optimisation of twist and rise

Although the local optimisation of helical twist and rise did converge to the correct solution in all four test cases, this is not necessarily the case in general. To provide insights into the robustness of the local optimisation of helical parameters, we performed multiple refinements with the MamK data set, starting from different initial estimates for the helical twist and rise (Supplementary Table 2). All these refinements were started from a cylindrical reference model that was low-pass filtered at 30 Å resolution. For convergence to the correct symmetry, values for the initial estimate of the twist need to lie within approximately ± 20% or 19~25° of the correct value; values for the initial estimates of the rise need to be within ± 20% or 42~62Å. Interestingly, relatively large errors in the initial estimates can still lead to correctly refined values, but in some cases the resolution of the corresponding refined map is suboptimal. This behaviour is likely due to the switching from exhaustive orientational searches in the initial stages of the refinement to local searches later on. This means that, if in a given refinement the parameters for twist and/or rise change considerably, one should probably perform a second refinement starting from the refined values of the first run. Indications for incorrectly refined symmetries are a lower resolution of the final map, and convergence of twist and/or rise to values at the edge of the user-defined search range.

### Pitfalls of imposing incorrect helical symmetry

To further illustrate the dangers of imposing an incorrect helical symmetry, we also re-refined the VipA/VipB data set using a twist of 64.9° and a rise of 3.63 Å. This symmetry corresponds to a 6-start helix with C1 symmetry, instead of the correct helix with C6 symmetry. After 3D auto-refinement and post-processing, RELION produces a map with a reported resolution of 4.5 Å (with refined values of twist=65.07° and rise=3.76 Å). Despite the relatively high nominal resolution, the map is devoid of protein-like features like alpha-helices or beta-strands that one would expect at this resolution (Fig 8A-B, left). In addition, the FSC curve shows "spiky" artefacts (Fig 8C). When the correct symmetry is imposed, the achieved resolution is 3.3 Å, the map does show the expected features (Fig 8A-B, right), and the FSC curve is smooth.

**Figure 8:**
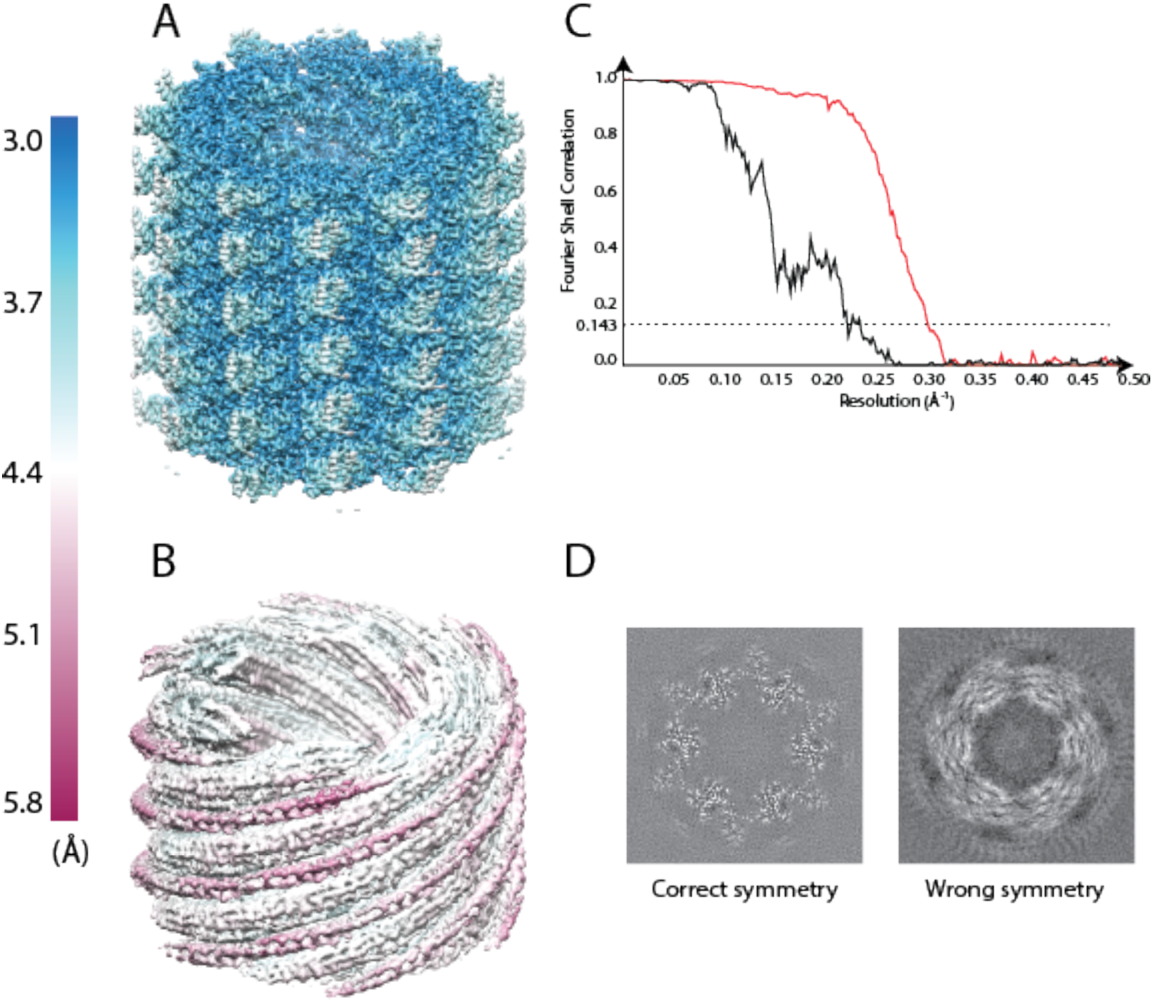
Effects of imposing different helical symmetries for the VipA/VipB data set. **A**. Reconstruction with the correct helical twist and rise imposed. B. Reconstruction with the wrong helical symmetry imposed. The maps in A and B are coloured according to local-resolution estimates (from 4 Å in blue to 6 Å in maroon). **C**. FSC curves between two independently refined half-sets for the refinement with the correct helical symmetry (red) and the wrong helical symmetry (black). **D**. Central slices through the unsharpened map from the refinement with the correct helical symmetry (left) and the wrong helical symmetry (right).

### The inter-box distance

To illustrate the effect of the inter-box distance on the reconstruction results, we performed refinement with various numbers of new asymmetric units per segment for TMV and for MAVS/CARD (Supplementary figure 2). For straight helical tubes like TMV, using a relatively large inter-box distance is expected to save computation time without affecting the final resolution. For more flexible helices like MAVS/CARD, shorter inter-box distance might be needed to describe the bends in the filaments with more orientational parameters. However, the results of our experiments suggest that larger inter-box distance still perform well, even for the more flexible MAVS/CARD filaments. In fact, in both cases inter-box distances in the range of 2-30% seem to yield structures of very similar quality. Possibly, segments corresponding to the most highly curved MAVS/CARD filaments are rejected by our auto-picking algorithm, or during the subsequent selection of 2D classes. An obvious advantage of using larger inter-box distances is that the computational costs of the refinement scale linearly with the total number of extracted segments.

### The half-sets for independent refinements

In 3D auto-refinement the data is divided into two half-sets, which are then refined independently in order to prevent an iterative built-up of noise due to overfitting (Scheres and Chen, 2012). For singleparticle analysis, the half-sets are created randomly from the extracted particles. For helical segments, random generation of the half-sets may be problematic, as the signal in one segment will partially overlap with the signal in its neighbouring segment. This overlap could introduce dependencies between the half-set refinements, and thus potentially lead to overfitting.

There are two aspects of our helical implementation that prevent this type of overfitting. Firstly, the prior on the translations (Fig 4A) only extends for a single rise in the direction of the helical axis. This will prevent the signal of any segment from translating on top of the identical, overlapping signal in the neighbouring segment. Secondly, to build in an additional safe-guard against overfitting, our implementation of helical processing divides the data set into half-sets based on a random division of helical filaments. That is, at the particle extraction stage, all segments from a single filament will be assigned to the same half-set.

To test both counter-overfitting measures, we performed 3D auto-refinements on the TMV data set (Supplementary Figure 3). In the first refinement, we used our implementation as intended, by both using the translational prior in the helical direction and the division of the data set into halves on the filament level. The post-processed FSC curve indicates a final resolution of 4.1 Å, which is in agreement with the features visible in the map. Importantly, beyond 3.8 Å, the FSC drops to values around zero for the rest of the frequencies. Secondly, we performed a refinement where we switched off both safe guards, i.e. we used an isotropic translational prior and random half-sets at the segment-level. In this case, severe overfitting of the data occurs as expected, as is evident from the FSC curve, which never drops to zero. A third refinement, where the half-sets were made at the filament-level but isotropic translational priors were used, shows no signs of overfitting. This refinement yielded a lower resolution map than the first refinement, probably as a result of a suboptimal translations along the helical axis. Finally, in a fourth refinement we made the half-sets at the segment-level and only used the anisotropic translational prior. In this case, a small amount of overfitting is present, indicating that the anisotropic translational prior alone is not enough to rigorously prevent overfitting.

### Angular and translational priors

In alternative refinement programs, the concept of priors on the rotations or translations are often expressed as local searches within a user-defined search range. Within the statistical framework, this corresponds to imposing a prior with a top-hat shape (i.e. with a constant, non-zero value within the search range, and a zero value outside the search range). In general, the better the shape of the prior reflects the actual distribution of the corresponding orientational parameters, the better the program is expected to perform, and the difference in performance is expected to become larger with increasing levels of noise in the data. Another advantage of the statistical framework, is that the optimal search range can be estimated from the data themselves, e.g. by optimising the variance in the translations perpendicular to the helical axis. A similar approach could in principle be implemented in the alternative approaches as well (Sigworth, 1998), but this is not done in many programs.

To test the influence of the shape of the prior on the refinement results, we compared standard refinements with Gaussian priors to refinements with top-hat priors. In the latter refinements, all tophat priors had non-zero values from −3 to +3 standard deviations of the corresponding Gaussian priors in the standard runs. Supplementary Figure 4 shows the resulting FSC curves after post-processing for these refinements on the MamK and VipA/VipB data sets. The difference between the Gaussian and the top-hat prior was not notable for MamK, but for VipA/VipB the Gaussian prior did yield a better structure.

## Discussion

We introduce functionality for helical reconstruction in the RELION program, and show for four different data sets that it can calculate maps to near-atomic resolution. For the three test cases from the EMPIAR data base, we show that helical processing in RELION yields results that are comparable to those obtained in alternative programs. Our implementation is integrated into the standardised workflow engine of RELION version 2.0, which facilitates the design of image processing strategy for novel users, and provides convenient tools for electronic bookkeeping and file management (Fernandez-Leiro and Scheres, 2016a). Because of the strong parallels with single particle analysis, users already familiar with RELION should find the transition to helical processing straightforward. The interaction with the user is similar to single-particle analysis of asymmetric particles, and parameters specific for helical reconstruction are mainly provided on dedicated tabs in the main GUI. Advantages that are inherent to the empirical Bayesian approach in RELION, such as the data-driven calculation of optimal filters for 3D reconstructions and alignment, are directly carried over from standard single-particle analysis to helical processing. In addition, the existing options for classification of images in 2D or in 3D, with or without masks, and with or without subtraction of partial signal (Bai et al., 2015; Scheres, 2016), are all available to be explored for helical processing of highly heterogeneous samples.

In the test cases presented in this paper, 2D classification proved useful in discarding suboptimal segments from the data set, but similar classification attempts in 3D were unsuccessful. These results contrast with common observations in single-particle analysis, where 3D classification is often key in selecting subsets of the data that yield the highest resolution maps (Fernandez-Leiro and Scheres, 2016b). On the one hand, these results may be attributed to the relatively high quality of the test data sets. On the other hand, there are intrinsic differences in classification of helices and individual asymmetric particles. The packing interactions inside the helical lattice, for example, may impose limitations on the structural heterogeneity of the individual protein subunits. And if structural heterogeneity does exist among the neighbouring subunits, for example because a floppy domain sticks out into the solvent, then classification of that heterogeneity may be difficult. The concept of a 3D helical class is particularly inadequate when the structural heterogeneity happens independently among individual segments along each filament. Only when different types of helical structures exist, i.e. when neighbouring segments in stretches of the filaments all adopt the same conformation, is 3D classification expected to perform well. For such cases, we implemented the option to independently refine the helical twist and rise of each class. Although this option did not improve the maps for the test cases described in this paper, we envision that it will be useful in cases where these types of structural heterogeneity do exist.

We explicitly stress that our implementation does not provide a generally applicable solution to the problem of local minima in the optimisation of helical parameters. Similar to alternative implementations of the IHRSR approach, RELION is based on an optimisation algorithm that will converge to the nearest local minimum, and the energy landscape of helical reconstruction, in particular with respect to the twist and rise parameters, is known to be highly complex (Egelman, 2010, 2014; Sachse et al., 2007). Thereby, the generation of a reliable initial 3D model with reasonable estimates for the helical twist and rise is likely to be the most difficult hurdle in helical reconstruction. For our test cases we employed two different methods. For MAVS/CARD and MamK, we successfully refined a featureless cylinder without imposing any point group or helical symmetry, and then estimated the helical twist and rise from the asymmetric reconstruction. This method is more likely to work for relatively simple helices, i.e. helices with few asymmetric subunits per box and few interactions between the subunits. An advantage of this method is that it has minimal user input. A drawback is that the initial few iterations of such refinements are relatively slow, as the probabilities for many orientations are significant, and these orientations thus all need to be considered in the reconstruction. For more complicated helices, asymmetric refinement of a featureless cylinder often does not work. For example, asymmetric refinement of a featureless cylinder never yielded a structure with discernable subunits for TMV or VipA/VipB. Fortunately, more complicated helices often yield more information-rich Fourier transforms, especially when these can be averaged together into high-resolution 2D class averages. Therefore, for complicated helices Fourier-Bessel indexing is a viable way to determine the initial estimate for helical twist and rise. For both TMV and VipA/VipB, placing spheres into the corresponding helical lattice then yielded an initial model within the radius of convergence of the refinement algorithm. Although computationally cheaper than refinement of a featureless cylinder, a drawback of this method is its dependence on user expertise in indexing diffraction patterns. Still, Fourier-Bessel indexing typically yields a better understanding of the structure, and may therefore help to avoid errors.

Our refinement of the VipA/VipB data set as a 6-start helix in C1 instead of the correct helical symmetry in C6 illustrates the pitfalls of imposing incorrect helical symmetry. Despite the incorrect symmetry, 3D auto-refinement yielded two half-maps that correlated with each other up to a resolution of 4.5 Å. Instead of directly measuring resolution, the FSC merely measures the consistency between two maps. By imposing the same, incorrect symmetry in both half-set refinements, the resulting maps become incorrect in a consistent manner, which results in a near-atomic resolution estimation of a completely incorrect map. At closer inspection, the reconstructed map did not reveal any protein-like features, and also the FSC curve itself showed artefacts. However, the indications that something is wrong may not necessarily be so clear. Actually, the MAVS/CARD test case was the subject of controversy itself, as an initial structure by the same authors was proven to be incorrect (Egelman, 2014; Xu et al., 2014, 2015). In our hands, starting MAVS/CARD refinements without imposing any symmetry from a featureless cylinder still led to the correct structure (Figure 6). Moreover, when starting from the incorrect helical parameters described in the original MAVS/CARD paper, the helical twist and rise consistently converged to the edge of the search ranges (not shown), which is a strong indication of incorrect estimates. Nevertheless, in the general case, validation of helical symmetry will remain difficult until one can reconstruct maps to near-atomic resolution, protein features like separated beta-strands and amino acid side chains can be readily distinguished, and a stereo-chemically sound atomic model can be refined inside the map. For cases where such maps cannot be obtained, doubts about the correct helical symmetry will remain relevant.

Apart from the initial model and the initial estimate for the helical twist and rise, important user-defined parameters are the inter-box distance and the different options to impose priors on the orientations. Smaller inter-box distances allow modelling more flexible filaments, but come at considerable computational cost, as each filament will yield more segments. Our results with TMV and MAVS/CARD indicate that relatively larger inter-box distances (i.e. up to 30%) may indeed be used without noticeable loss in resolution. When in doubt, one could start processing with a relatively large inter-box distance, and then re-run refinements near the end with a smaller inter-box distance to make sure the best possible resolution is obtained. Our results with VipA/VipB indicate that Gaussian-shaped priors may add useful information to the refinement when compared to top-hat priors, which are typically employed in alternative programs. This difference however was not noticeable in the case of MamK. Still, because these priors come at virtually no extra computational cost and Gaussian distributions are more likely to describe the orientational distributions in experimental data, the Gaussian priors in RELION are switched on by default.

Within the field of single-particle analysis, considerable concern has arisen about the accurate estimation of resolution in 3D reconstructions (Henderson et al., 2012). One approach to prevent the iterative build-up of high-resolution noise in the reconstruction is the independent refinement of two halves of the data (Scheres and Chen, 2012). In standard single-particle analysis, the two half sets are often generated randomly or based on odd/even particle numbers. When extracting overlapping segments from helical filaments, the overlap in signal between neighbouring segments could introduce spurious dependencies between the two half-sets. Our results show that a random division of the data set based on filaments rather than segments, combined with a tophat prior on the translations along the helical axis, effectively yield maps with reliable resolution estimates.

In conclusion, we present an implementation of the IHRSR approach to helical processing in the RELION program. The object-oriented character of RELION's code facilitates the re-use of many of its functions. This makes it relatively straightforward to extend the helical processing approach presented here to 3D data, such as sub-tomograms. This extension is something we will pursue for a future version. Another possible extension is the introduction of functionality to deal with breaks in the helical symmetry, such as the seam in microtubules (Metlagel et al., 2007). The code available in the current 2.0 version implements all features described in this paper, and has already been useful in calculating several new structures in our own lab. We hope this paper will help to make it similarly useful to others.

## Acknowledgements

We thank Nigel Unwin, Tony Crowther, Jude Short and Jan Löwe for stimulating discussions, and Jake Grimmett and Toby Darling for assistance with high-performance computing. This work was funded by the UK Medical Research Council (MC_UP_A025_1013).

